# Removing unwanted variation in a differential methylation analysis of Illumina HumanMethylation450 array data

**DOI:** 10.1101/019042

**Authors:** Jovana Maksimovic, Johann A. Gagnon-Bartsch, Terence P. Speed, Alicia Oshlack

## Abstract

Due to their relatively low-cost per sample and broad, gene-centric coverage of CpGs across the human genome, Illumina’s 450k arrays are widely used in large scale differential methylation studies. However, by their very nature, large studies are particularly susceptible to the effects of unwanted variation. The effects of unwanted variation have been extensively documented in gene expression array studies and numerous methods have been developed to mitigate these effects. However, there has been much less research focused on the appropriate methodology to use for accounting for unwanted variation in methylation array studies. Here we present a novel 2-stage approach using RUV-inverse in a differential methylation analysis of 450k data and show that it outperforms existing methods.

## INTRODUCTION

DNA methylation, which is the addition of a methyl (CH_3_) group to the cytosine of a CpG dinucleotide, is the most widely studied epigenetic modification in human development (1) and disease (2–4). As interest in epigenetics has grown, Illumina’s Infinium HumanMethylation450 (450k) arrays have emerged as a popular platform for genome-wide methylation analysis, particularly for projects requiring large numbers of samples. Its broad coverage of the human genome (>450,000 CpGs) and relatively low cost per sample has resulted in the extensive use of 450k methylation arrays in several large studies such as The Cancer Genome Atlas (TCGA), Encyclopaedia of DNA Elements (ENCODE), and numerous Epigenome-Wide Association Studies (EWAS)(5–7).

Unfortunately, large studies can be particularly susceptible to the effects of unwanted technical variation due to the large number of samples requiring processing. For example, processing may have to occur over several days or be performed by multiple researchers thus increasing the likelihood of technical differences between “batches”. Furthermore, unwanted technical variation is often present against a background of unwanted biological variation. For example, EWAS are often performed using blood as it is an easily accessible tissue; however, blood is a heterogeneous collection of various cell types, each with a distinct DNA methylation profile. Many recent studies have highlighted the need to account for cell composition when analysing DNA methylation (8–10) as it has been shown to influence differential methylation (DM) calls (6, 11–15).

The impact of unwanted variation such as batch effects, has been extensively documented in the literature on gene expression microarrays (16, 17) and numerous methods have been developed for correcting for unwanted variation in expression array studies. When the sources of unwanted variation are “known”, it is common to incorporate an additional factor into a linear model to explicitly account for batch effects, or to apply a method such as ComBat, which uses an empirical Bayes (EB) framework to adjust for “known” batches (18). However, sometimes the source(s) of unwanted variation are unknown. For example, a sample of sorted cells may contain contaminating cells of another type and the level of contamination may vary between samples. This introduces unwanted variation into the data, however the source of the variation may not be obvious and is thus impossible to model. In such cases, methods such as Surrogate Variable Analysis (SVA) (19, 20) and Independent Surrogate Variable Analysis (ISVA)(21) attempt to infer the unwanted variation from the data itself. Recently, Gagnon-Bartsch and Speed (22) published a new method, Remove Unwanted Variation, 2-Step (RUV-2), which introduced the concept of estimating the unwanted variation using negative control features that should not be associated with the factor of interest but are affected by the unwanted variation. More recently, the authors have extended their work on RUV-2 to develop RUV-inverse and several other variations (23).

RUV-2 uses factor analysis of the negative control features to estimate the components of unwanted variation. A number, *k*, of the unwanted factors are then included in a linear model to perform the adjustment. The choice of *k* is critical to the performance of the algorithm but there is no straightforward way to select *k* (22). RUV-inverse removes the need to empirically determine the “best” *k* and, unlike RUV-2, is also relatively robust to the misspecification of negative control features (23).

RUV-2 has been successfully applied to metabolomics, gene expression and 450k methylation array data (8, 22, 24). Compared to RUV-2, RUV-inverse has shown improved performance on gene expression data (23). Given that RUV-inverse offers both usability and performance improvements over RUV-2 (23) it could prove useful in mitigating the effects of unwanted variation in 450k array studies. However, as different data types have different properties, it is not obvious how to apply the method to 450k data to obtain the best results. For example, 450k arrays contain over 450,000 features as opposed to the ∼20,000 present on gene expression arrays and there is no direct analogue of house-keeping genes in the methylation context. As such we have developed a novel, 2-stage approach specific to using RUV-inverse with 450k methylation data (Figure 1).

**Figure 1.**
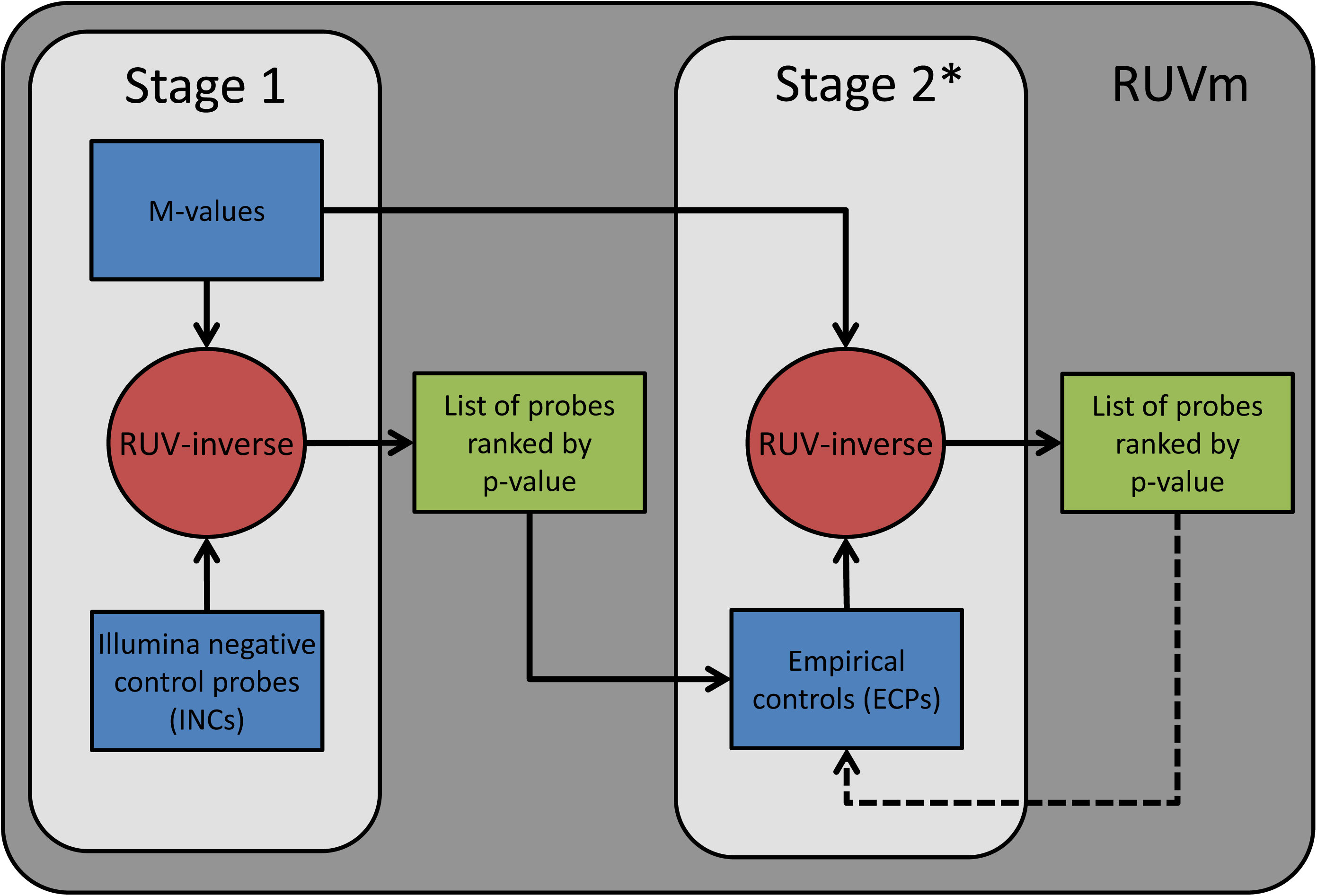
A schematic representation of a DM analysis using RUVm. The RUVm approach has two stages. The red circles indicate a DM analysis step. The blue rectangles represent the inputs that are required for each stage. The green rectangles are the outputs that are produced by each stage. Stage 1: Perform an initial DM analysis using RUV-inverse with Illumina negative control probes (INCs); CpGs are ranked by p-value based on the strength of their association with the factor of interest. Stage 2: Use the list of CpGs from Stage 1 to select a set of ECPs which are then used in a DM analysis with RUV-inverse. (*) Stage 2 can be performed one or more times.

The ability to robustly correct for unwanted variation in 450k methylation array data would not only aid in improving the results of individual studies, it would also enable the effective integration of data on the same samples from different studies/sources, resulting in increased statistical power to detect true DM. Linear regression with adjustment, ComBat (18), SVA (19, 20) and ISVA (21) have all been applied to different types of methylation array data in various contexts (21, 25–29), however there has not been a comprehensive assessment of the relative performance of *all* these methods in a single study, focused on 450k data. Here we present our 2-stage approach, RUVm, for the application of RUV-inverse to 450k methylation data and evaluate it against the methods previously outlined. We show that our approach is robust and consistent and often outperforms other methods in a differential analysis of 450k data. We make the method for 450k analysis freely available in the *missMethyl* Bioconductor package.

## MATERIAL AND METHODS

### Data processing and analysis

All 450k data was imported into the R statistical computing environment (3.1.0) (30) using the Bioconductor (3.0) package *minfi* (1.12.0) (31). Data was filtered based on the following criteria: poor quality samples (mean detection P-value > 0.01) were discarded; probes with a detection p-value > 0.01 in at least 1 sample within a dataset were also discarded, as were X and Y chromosome probes, probes with SNPs at the CpG or single base extension site and the cross-reactive probes identified by Chen et al. (32).

All DM analyses were performed on M-values (M = log_2_(methylated/unmethylated)) as recommended by Du et al. (33). Regression analysis was performed using the empirical Bayes methodology available in *limma* (3.22.1) (34, 35). We used the reference implementations of the ComBat (36) and SVA (19, 20) methods in the *sva* package (3.12.0) (37). Known batches were passed as a variable to the ComBat function and we allowed SVA to estimate all the surrogate variables. Both methods were used in conjunction with *limma*. We also used the reference implementation of ISVA (21) from the *isva* CRAN package. As with SVA, we allowed ISVA to estimate all the surrogate variables. RUVm analysis was performed using Bioconductor *missMethyl* (1.1.1) package implementation, with method = “inv”. Unless otherwise stated, all p-values reported were adjusted for false discovery rate using the Benjamini-Hochberg method (38). The R code for all the analyses is available in Additional Files 2-6.

### RUV-inverse

With RUV-inverse, differential methylation is estimated using a generalized least squares (GLS) regression. The covariance matrix that is used is the empirical covariance matrix of the negative controls. Some difficulty arises in the calculation of the standard errors. The “traditional” GLS standard errors end up being the same for every CpG; however, this is undesirable, as we believe that some CpGs are more variable than others. To solve this problem, and allow different CpGs to have different standard errors, we calculate the standard errors using the “inverse method,” described in detail in Gagnon-Bartsch et al. (23), and also summarized briefly in Additional File 1.

The basic idea of the inverse method is to re-fit the model, but including an extra, randomly-generated column in the design matrix. The estimated regression coefficient for this random column should be about zero, because there is no “true effect” associated with this random column. The extent to which the estimated regression coefficient is *not* precisely equal to zero for any particular CpG gives us information about the variability of that CpG. This information can be used (after repeating the procedure many times) to calculate a standard error for the CpG.

### Ageing data

#### Test set

The birth versus 1.5 years (Study 1) methylation data was published by Martino et al. (39) (GSE42700). Briefly, buccal cells were collected from 30 individuals at birth and 1.5 years. The cohort included 10 monozygotic twin pairs and 5 dizygotic twin pairs.

The birth versus 1 year (Study 2) methylation data was published by Martino et al. (40) (GSE34639). Blood samples were collected from 48 individuals at birth and again at 1 year of age. Half of the samples from each time point were cultured in either standard media or media containing anti-CD3 antibody; CD4+ T-cells were then isolated by positive selection. DNA from 2 individuals was pooled into a single sample at each time point, resulting in 12 conventional and 12 anti-CD3 birth samples and 12 conventional and 12 anti-CD3 1 year samples. As the authors did not find any significant methylation differences between the conventional and anti-CD3 treated samples we elected to only utilise the conventional samples in this study, leaving 12 birth and 12 1 year samples.

The birth versus 100 year (Study 3) methylation data was published by Heyn et al. (41) (GSE30870). Peripheral blood was taken from 20 healthy centenarian donors and umbilical cord blood was taken from 20 newborns. For 19 of the newborn and centenarian samples, DNA was extracted from either CBMCs or PBMCs, respectively. DNA was extracted from CD4+ T cells purified from the remaining birth and 100 year sample. We only used data from the 38 unsorted samples in this study.

The 30 Study 1 samples, 24 Study 2 samples and 38 Study 3 samples were then combined into a single dataset and filtered as previously described, leaving 401,057 probes. The combined dataset was also pre-processed using 2 methods: SWAN (42) and Stratified Quantile Normalisation (SQN) (43), producing 2 different datasets. The sample identifiers and descriptions can be found in Supplementary Table 1.

#### Truth set

The birth versus 18 years data was published by Cruickshank et al. (44) (GSE51180). The study was comprised of 24 subjects; 12 who were born prematurely and 12 who were born at term. DNA was extracted for each subject at birth from a neonatal Guthrie card and at 18 years of age from a dried blood spot; resulting in a total of 48 DNA samples. One of the term birth samples was excluded by the authors as it failed quality control, leaving 47 samples.

The data was pre-processed using SWAN (42) and filtered as previously described, leaving 395,173 CpGs. Blood cell type proportions were estimated using the ‘estimateCellCounts’ *minfi* function, which implements the method described by Jaffe and Irizarry (8). The regression analysis for detecting DM between birth and 18 years was performed using *limma*; to adjust for differences in cell type composition between the ages, the cell type proportions previously estimated using ‘estimateCellCounts’ were included as covariates in the linear model.

### Smoking data

#### Test set

The smoking methylation data was originally published by Liu et al. (6)(GSE42861) as part of their study examining the association between methylation and rheumatoid arthritis. We used the 200 current smoker and 193 never smoker samples in our analysis (Supplementary Table 2). These included a mix of rheumatoid arthritis patients and controls of both sexes and a range of ages. The DNA was extracted from EDTA-treated blood.

An additional 656 methylation samples with unknown smoking status were obtained from the dataset originally published by Hannum et al. (45) (GSE40279). DNA was extracted from whole blood samples collected from both male and female individuals, of two different ethnicities and a range of ages. In this study, we used 70 randomly selected samples from the Hannum data and combined them with a random sample of 80 current smokers and 50 never smokers from the Liu data to create a dataset with significant unwanted variation (Supplementary Table 3). We generated another two datasets using the same approach but with different combinations of randomly selected samples (Supplementary Table 4, 5).

#### Truth set

We used the 187 CpGs smoking-associated CpGs published by Zeilinger et al. (46) as the “truth” set for our analysis (Supplementary Table 6). These CpGs were originally identified by the authors in a discovery cohort of 262 current smokers and 749 never smokers and were subsequently replicated in a second cohort of 236 current smokers and 232 never smokers (46). The DNA was extracted from whole blood.

### Cancer data

#### Test set

The Lung Adenocarcinoma (LUAD) data, which is comprised of 427 tumour and 31 normal samples, was obtained from TCGA. All data was downloaded from the TCGA Data Portal as unprocessed IDAT files. In our analysis, we used all 31 normal samples and 75 tumour samples that were on the same BeadChips as the normal samples (Supplementary Table 7). The data was pre-processed using both SWAN (42) and *minfi* functional normalisation (FNORM) (47), producing 2 different datasets. Each of the pre-processed datasets was filtered as described previously, leaving 411,735 CpGs.

#### Truth set

The LUAD bisulfite sequencing data was published by Zheng et al. (48) (GSE56712). Five lung tumour and 5 matched normal samples were taken from 5 patients with LUAD. DNA was extracted and captured using the Agilent SureSelect Methyl-Seq system, followed by bisulfite sequencing, resulting in 15-40 million 90bp paired-end Illumina reads per sample (48).

We obtained the raw FASTQ files from the Sequence Read Archive (SRA). The reads were assessed for quality using FastQC (0.10.1). Trimming was performed with Trim Galore (0.3.7); 10bp were trimmed from the 5’ end of each read and 5bp were trimmed from the 3’ end of each read following quality trimming and adapter removal. Read pairs were discarded if at least one read from the pair was less than 20bp long. The reads were then mapped to the human genome (hg19) using Bismark (0.12.5) and Bowtie2 (2.1.0). Duplicates were removed with the deduplicate_bismark tool. Methylation calls were made using the bismark_methylation_extractor. The pipeline for processing the bisulfite sequencing data was implemented in bpipe (0.9.8.6) (49) and is available at https://github.com/JovMaksimovic/methyl-seq_bpipe. The data was then imported into R (3.1.0) for downstream analysis. CpGs not covered by at least 1 read in *all* 10 sequenced samples were discarded, leaving 1,656,501 loci per sample. We then identified the CpGs that were covered in *both* the filtered LUAD 450k dataset and the bisulfite sequencing dataset, which left 221,694 CpGs for downstream analysis. Differential methylation analysis of the 5 tumour versus 5 normal methyl-seq samples was performed using the Bayesian hierarchical model and Wald test approach implemented in the Bioconductor *DSS* package (50).

## RESULTS

### RUVm: A 2-stage approach for differential methylation analysis of 450k data using RUV-inverse

As with RUV-2, performing a DM analysis using RUV-inverse relies on having negative control features to accurately estimate the components of unwanted variation (see Methods)(22, 23). Gagnon-Bartsch and Speed (22) emphasise that the choice of negative control features can be crucial in determining the effectiveness of the method. Negative control features are probes/genes that are known *a priori* not to be associated with the biological factor of interest, but are affected by unwanted variation. For example, in a microarray gene expression study, these could be house-keeping genes or a set of spike-in controls (22). Although there is some evidence that the CpGs in the CpG islands of housekeeping gene promoters are generally unmethylated (51), to our knowledge, there is no general list of “housekeeping” CpGs for researchers to draw from that could be used as negative control features in a differential methylation analysis with RUV-inverse.

Given that CpG methylation varies greatly between different cell types, tissues etc. (52, 53), it would be beneficial to be able to empirically determine CpG probes that are not associated with a particular factor of interest for each individual experiment. However, empirically identifying negative control probes shares the same difficulties inherent to determining which probes are differentially methylated, particularly in the presence of unwanted variation. Gagnon-Bartsch and Speed (22, 23) outlined a strategy in which an initial DM analysis is used to determine empirical control probes (ECPs) for use in a subsequent DM analysis. A similar iterated model was also previously proposed by Leek and Storey (20). The basic requirements for this strategy to work are that the initial DM analysis is “good enough” and that the subsequent DM analysis is somewhat robust to an imperfect set of negative control features.

Based on these criteria, we propose using a 2-stage approach for the DM analysis of 450k data (Figure 1). Stage 1 involves performing a DM analysis using RUV-inverse with the 613 Illumina negative controls (INCs) present on the 450k array to rank all the CpG probes by p-value based on their association with the factor of interest. The INCs are randomly permuted sequences that should not hybridize to the DNA template and are generally used to define the system background. Thus, they are not expected to contain any biological signal but do capture some technical variation between samples, chips, batches etc. which can result in an improvement in probe rankings over an unadjusted analysis. However, as the INCs only produce a background level signal and can only capture technical variation they are not an ideal set of negative control features. Hence, our approach uses the results from Stage 1 to empirically select a more informative set of negative controls from the CpG probes on the 450k array. This involves designating a proportion of the least associated CpG probes as ECPs for use in Stage 2. In Stage 2, the ECPs are used to perform a second DM analysis of the original dataset with RUV-inverse. If necessary, Stage 2 can be performed multiple times to further refine the set of ECPs, although this often not necessary (23). For simplicity, the 2-stage approach described will henceforth be referred to as “RUVm”.

### Ageing methylation data

Batch effects occur because measurements can be affected by factors such as laboratory conditions, differences in reagents and/or equipment or because different personnel processed different samples (16, 17). To determine how various methods perform at correcting for large batch effects in 450k methylation data, we created a dataset with a pronounced batch effect by merging data from three different studies concerned with examining changes in methylation due to age. The three contributing studies (Study 1-3) all compared methylation at birth to methylation in older individuals (see Methods for details). In the resulting dataset, which will henceforth be referred to as the “ageing+” data, the batch effects are, in fact, much larger than the factor of interest (Figure 2c-f). This is analogous to many EWAS in which the collection and processing of numerous samples makes them susceptible to batch effects that are much larger than the effect of interest (case versus control).

**Figure 2.**
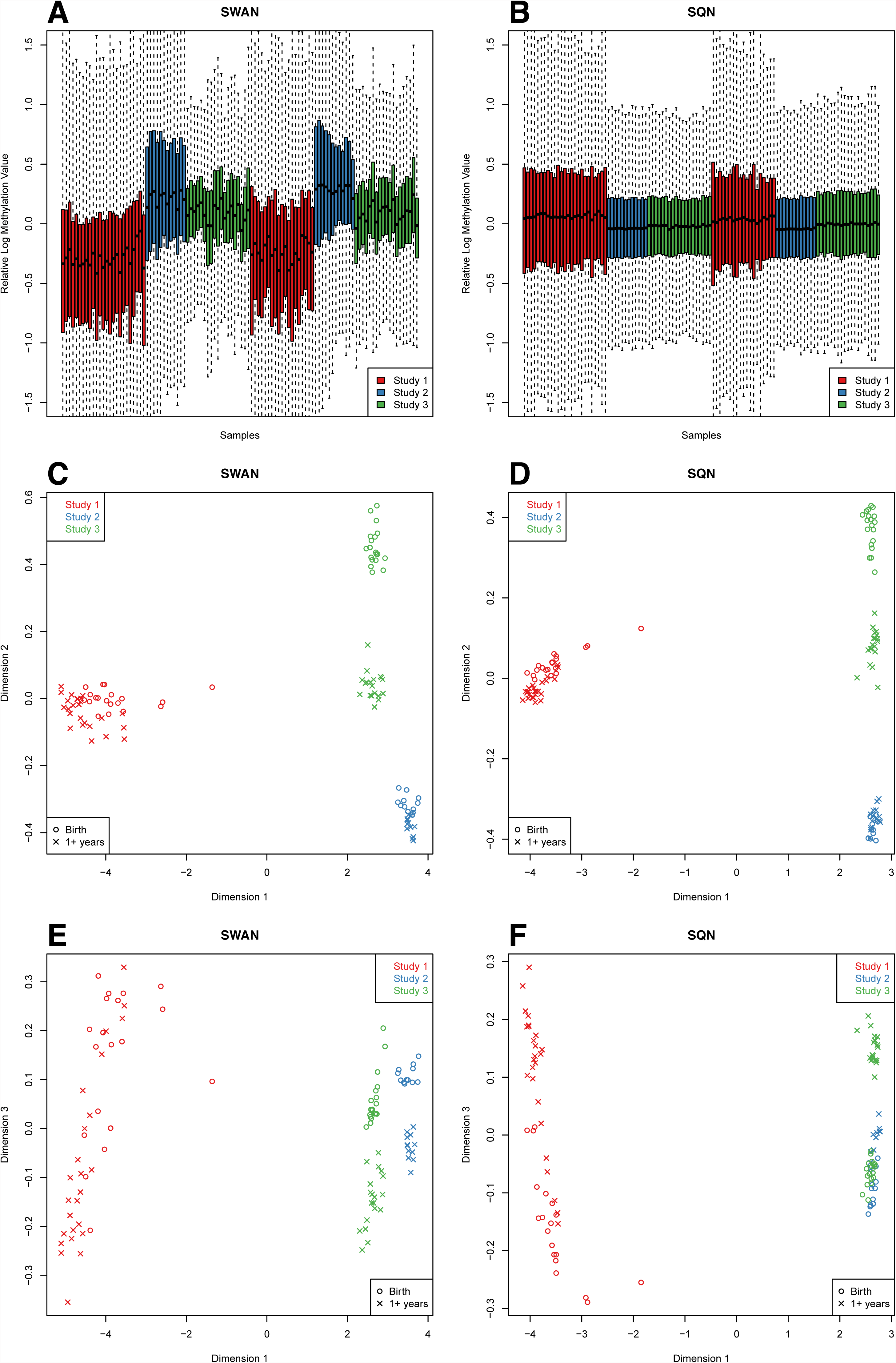
Relative Log Expression (RLE) and Multi-Dimensional Scaling (MDS) plots of the ageing+ data. These RLE plots show the deviation from the median methylation level (M-value) for each of the 450k arrays from the 3 ageing studies combined. An MDS plot is analogous to a principal components analysis plot. The axes represent the major sources of variation in the data based on the top 1000 genes with the largest standard deviations between samples; dimension 1 represents the largest source of variation, dimension 2 represents the next largest orthogonal source of variation, followed by dimension 3, etc. (A) RLE plot: SWAN pre-processed data. The samples are coloured by the study the data originated from: Study 1, 2 or 3. (B) RLE plot: SQN pre-processed data. The samples are coloured by the study the data originated from: Study 1, 2 or 3. (C) The MDS plot of the first 2 dimensions of the SWAN pre-processed data shows that the largest source of variation between samples is tissue type. (D) The MDS plot of the first 2 dimensions of the SQN pre-processed data shows that the largest source of variation between samples is tissue type. (E) The MDS plot of dimension 3 versus dimension 1 of the SWAN pre-processed data shows that age is the third largest source of variation in the data. (F) The MDS plot of dimension 3 versus dimension 1 of the SQN pre-processed data shows that age is the third largest source of variation in the data.

In gene expression studies, relative log expression (RLE) plots are commonly used to show deviation from the median gene expression level, to determine the overall quality of the dataset and to identify poor arrays. In the case of 450k data, we look at the deviation from the median methylation level for each array on the M-value scale. The RLE plot in Figure 2a highlights the existence of 3 clear batches in the ageing+ data with the application SWAN (42) within-array pre-processing. Using SQN pre-processing (43), which normalises between arrays as well as between probe types, improves the appearance of the RLE plot (Figure 2b), however there are still differences between the 3 studies, particularly Study 1 compared to 2 and 3, indicating that the batch effects have not been eliminated.

Multi-dimensional scaling (MDS) plots of the data using the 1000 most variable probes show that the largest source of variation is cell type, regardless of the type of pre-processing used (Figure 2c, d). Unsurprisingly, the buccal cell samples from Study 1 are distinct from the blood-derived samples from Studies 2 and 3. Furthermore, the Study 2 samples, which were extracted from purified CD4+ T-cells, show significantly less variability than the Study 3 whole blood samples. Both the birth and 1 year Study 2 samples cluster closer to the Study 3 birth samples than to the Study 3 centenarian samples. Examining a higher dimension reveals that age is associated with the third largest dimension of variation in the data (Figure 2e, f). Figure 2f also shows that using SQN (54) pre-processing reduces the variation between Studies 2 and 3 compared to using SWAN (42)(Figure 2e).

#### Defining the ageing “truth” set

To compare the relative performance of methods for removing batch effects in methylation array data, an appropriate “truth” dataset was required. We used an unrelated dataset, published by Cruickshank et al. (44), which measured methylation in blood at birth and 18 years of age in a cohort of 24 individuals to identify CpGs that change in methylation with age (see Methods). However, as Jaffe and Irizarry (8) recently demonstrated, changes in DNA methylation can be confounded with changes in cell type proportions when methylation is measured in a mixed cell population, such as blood. Consequently, as Cruickshank et al. (44) data was derived from Guthrie cards and dried blood spots, we needed to ensure that the “truth” set was not contaminated with spurious associations due to changes in cell type proportions confounded with age.

Firstly, we sought to characterise whether there was significant confounding between the age at which the samples were taken and changes in blood cell type composition between birth and 18 years. We used the method developed by Jaffe and Irizarry (8) to estimate the proportions of the various blood cell sub types in the birth and 18 year old samples. Supplementary Figure 1a shows that although there were some differences in cell type proportions that correlated with age, the confounding is not extreme. In their study, Jaffe and Irizarry (8) showed that in the Heyn et al. (55) birth versus centenarian data, the centenarian samples were almost entirely comprised of granulocytes, meaning that a comparison of methylation between ages in that dataset is effectively a comparison between cell types. We recapitulated their results in this study (Supplementary Figure 1b) to illustrate that the Cruickshank et al. (44) data was not as dramatically affected by changes in cell type proportions. Furthermore, examining the methylation distribution of the 600 probes identified by Jaffe and Irizarry (8) as discriminating between blood cell sub types (Supplementary Figure 2), indicates that there is no significant difference in the distribution of these probes between birth and 18 years (K-S Test p-value = 0.29) in the Cruickshank et al. (44) data (Supplementary Figure 1c). In contrast, the difference is statistically significant for the Heyn et al. data (K-S Test p-value < 2 ×10^-16^) (Supplementary Figure 1d), due to the large difference in cell type proportions between the newborn and centenarian samples. It is also apparent that the methylation distribution of the probes from the centenarian samples most closely resembles that of the granulocytes, which further supports the proportion estimate shown in Supplementary Figure 1b. Finally, a DM analysis using only the 600 cell type discriminating probes resulted in 268 significant associations (FDR adj. p-value < 0.05) with age for the Cruickshank et al. data and 506 for the Heyn et al. (Supplementary Figure 1e).

Despite demonstrating that the Cruickshank et al. (44) data does not show dramatic differences in estimated cell type proportions between the birth and 18 years samples, differences do exist that may contribute to false associations between methylation and age. To mitigate these effects we included the estimated cell type proportions as covariates in the *limma* linear model used to identify differential methylation between birth and 18 years. Prior to adjusting for cell type proportions we identified 100,800 significantly DM CpGs at FDR adj. p-value < 0.05. However, after including the cell type proportion estimates in the linear model, our analysis ultimately identified 2,238 “true” positives (FDR adj. p-value < 0.05) and 188,895 “true” negatives (FDR adj. p-value > 0.9).

#### Differential methylation analysis of ageing+ data

Before comparing the performance of RUVm to that of other methods, we were interested in evaluating the effect of ECP selection on the performance of RUVm. The selection of ECPs must be based on some criteria for determining the proportion of probes that are the “least” associated with the factor of interest. One possibility is to choose ECPs based on a cut-off using FDR adjusted p-values (23). This will often work, but may fail if the p-values are biased (either systematically inflated or deflated). Another option is to select a fixed percentage of the lowest ranked probes based on expected amount of DM in the data; for example, it is expected that a DM study of cancer versus normal would identify many DM probes whereas an EWAS investigating the effect of maternal smoking during pregnancy is expected to result in very few DM loci. To evaluate how selection of ECPs affected the performance of RUVm, we performed several analyses on the ageing+ data, which was pre-processed in 2 different ways (SWAN and SQN) and varied how ECPs were selected (FDR > 0.1-0.9 or bottom 10-90%). We gauged relative performance by constructing (Receiver Operating Characteristic) ROC curves using the truth data described in the previous section.

Supplementary Figure 3 shows that the proportion of probes selected as ECPs decreases linearly with increasing FDR cut-off in the range of 0.1-0.9. It also shows that pre-processing affects the proportion of probes selected as ECPs by FDR cut-off, which is unsurprising as normalization techniques are designed to reduce the variability between samples and thus influence p-values (56– 58). Supplementary Figure 4 shows that, for the ageing+ data, the performance of RUVm is relatively robust to the choice of ECPs. This result is consistent irrespective of pre-processing or the criteria used to select ECPs: either FDR cut-off or fixed percentage.

**Figure 3.**
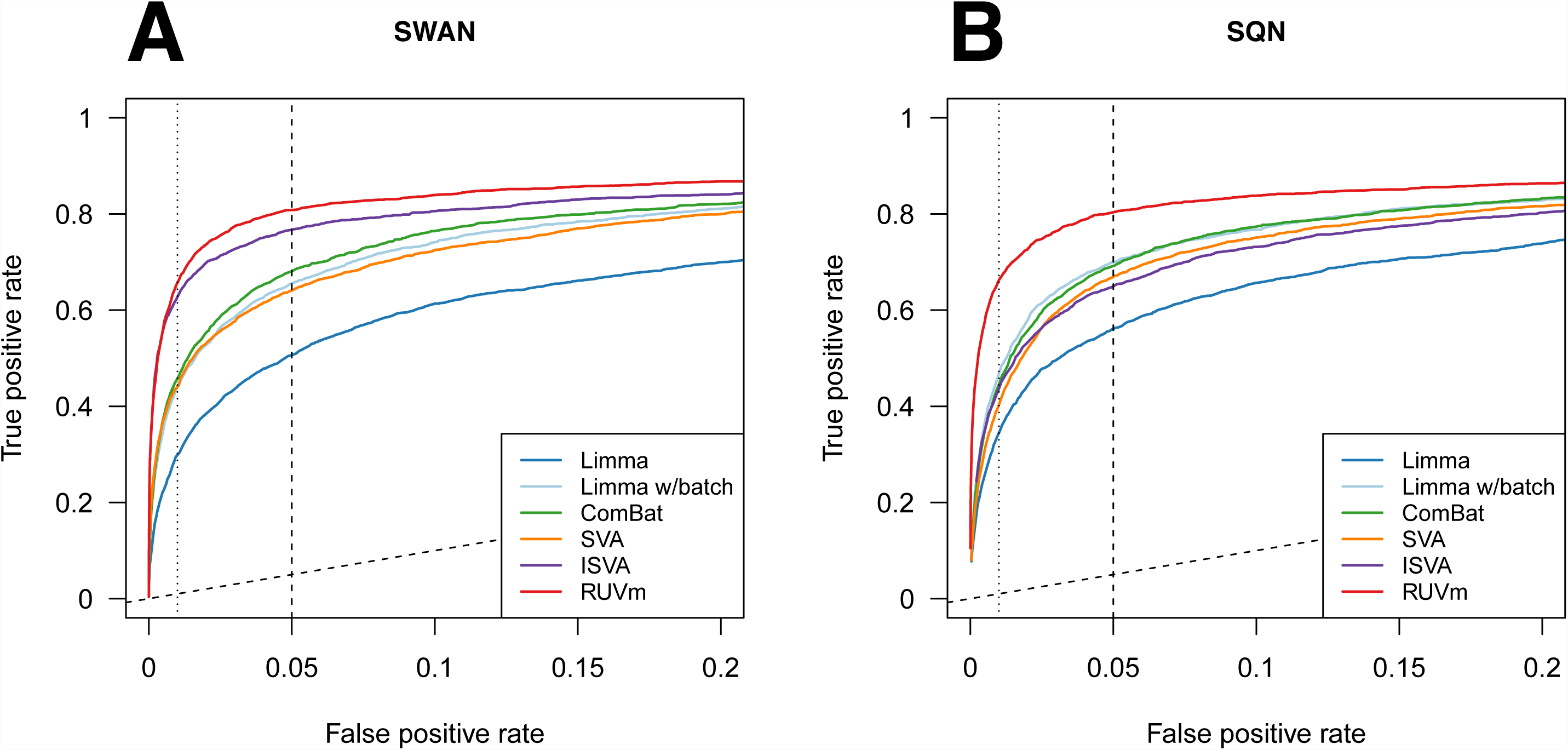
Performance of various adjustment methods in a DM analysis of the ageing+ data. (A) ROC curve showing the false positive rate versus the true positive rate for the various adjustment methods on the SWAN pre-processed data. (B) ROC curve showing the false positive rate versus the true positive rate for the various adjustment methods on the SQN pre-processed data.

**Figure 4.**
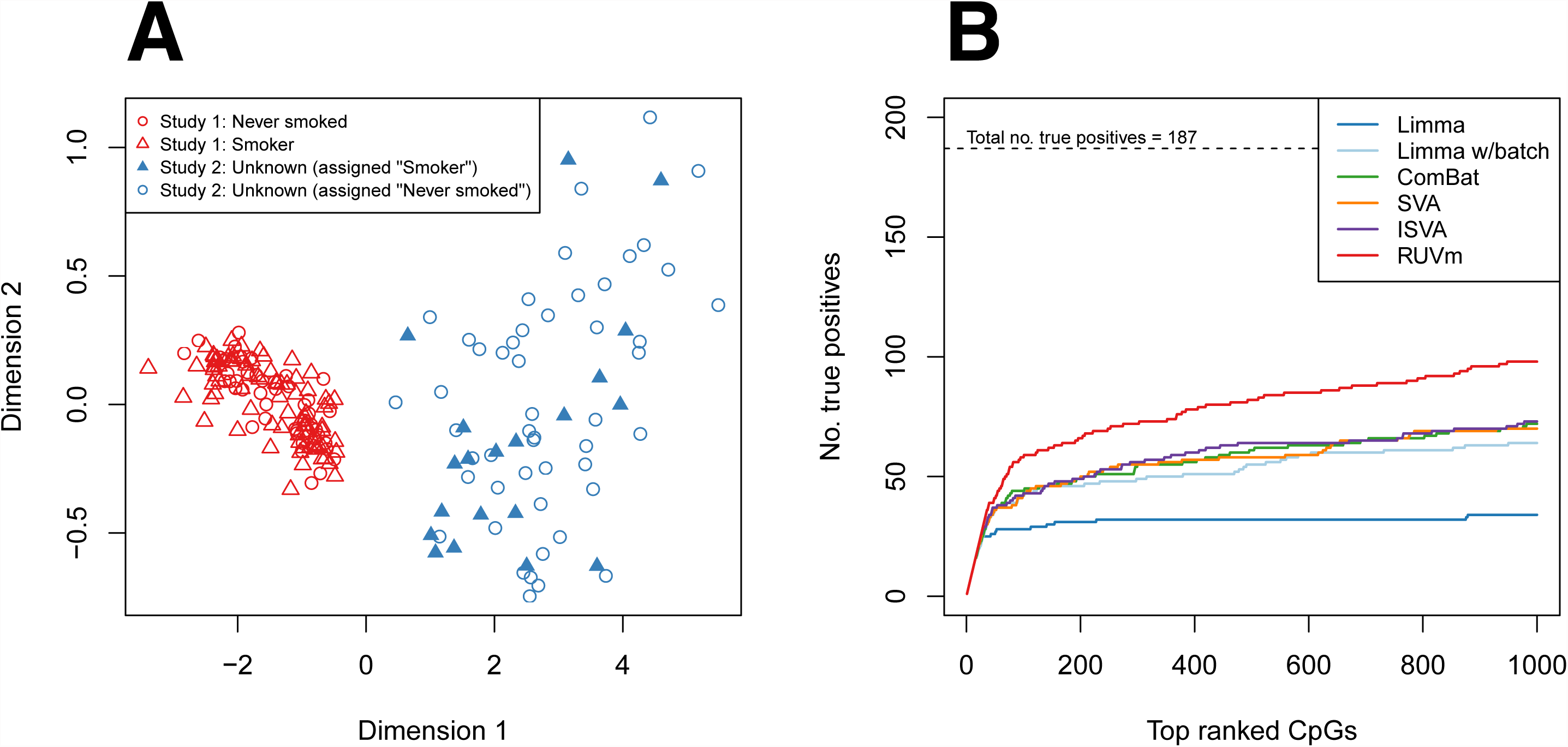
DM analysis of the smoking+ data. (A) MDS plot of the first 2 dimensions of the SWAN pre-processed smoking+ data shows that the largest source of variation between samples is the study of origin and there is no visible clustering by smoking status. The smoking+ data consists of a combination of 80 smokers and 50 never smokers from the Liu data and 70 samples from the Hannum data, 20 of which were assigned as “smokers” and 50 as “never” smokers. (B) Performance of various analysis methods in a DM analysis of the smoking+ data. The lines represent the cumulative number of true positives ranked in the top 1000 CpGs produced by the different methods.

We then compared the performance of RUVm to several other methods in a DM analysis of the ageing+ data; an unadjusted *limma* regression analysis, an “adjusted” *limma* regression analysis including a factor for study, ComBat (18) with study as the batch variable, ISVA (21) and SVA (19, 20). We used the 2-stage RUVm approach previously described with ECPs selected at FDR adjusted p-value > 0.2. Evaluating the quality of an adjustment on real data is not trivial. Gagnon-Bartsch and Speed (22) suggest several useful strategies such as looking at p-value distributions (19, 20) and the rankings of “true” positives. If “true” negatives are also available, an ROC curve can be particularly useful as it allows the visualisation of the rate of false positives versus the rate of true positives.

The p-value histograms resulting from the DM analyses of the ageing+ data using various approaches are not a dramatic departure from the ideal shape, which is expected to be almost uniform with a spike near zero (Supplementary Figure 5). However, regardless of the type of pre-processing used, the p-value histogram for the *limma* analysis without the batch factor does have a significantly shorter bar and the most pronounced slope in the height of the bars between 0.05 and 1 (Supplementary Figure 5a, b). This suggests that, as expected, the “batch adjusted” analysis is an improvement for *limma*. However, as the p-value histograms for the other methods look very similar, using p-value histograms alone is insufficient to discern the relative performance of the other algorithms.

**Figure 5.**
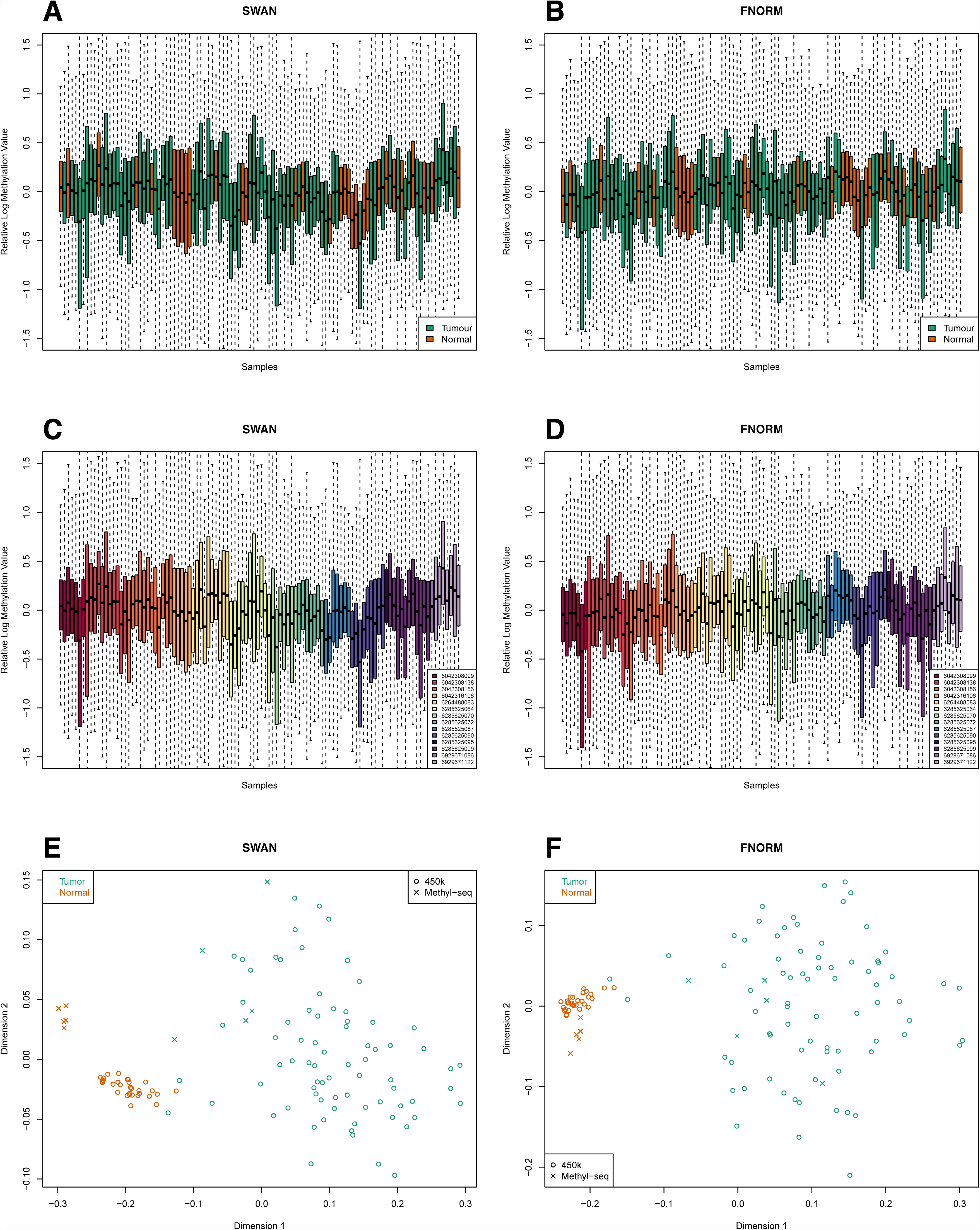
RLE and MDS plots of a subset of the TCGA LUAD 450k methylation data; includes all 31 normal samples and 75 of the tumour samples that shared chips with the normal samples. (A) SWAN pre-processed data; coloured by tumour/normal status. (B) FNORM pre-processed data; coloured by tumour/normal status. (C) SWAN pre-processed data; coloured by chip. (D) FNORM pre-processed data; coloured by chip. (E) MDS plot of both the TCGA LUAD data and the methyl-seq LUAD data. Uses the top 1000 most variable CpGs of the 221,694 loci covered by both platforms after filtering; 450k data pre-processed using SWAN. (F) As for (E); 450k data pre-processed using FNORM.

An ROC analysis using the Cruickshank et al. (44) “truth” data shows that RUVm outperforms other methods regardless of the type of pre-processing used (Figure 3). When the data was pre-processed with SWAN (42), RUVm performed the best, followed closely by ISVA. ComBat was the next best performing method, although its performance was substantially lower than that of ISVA, followed by *limma* with a factor for study and SVA. Unsurprisingly, the unadjusted *limma* analysis performed the poorest (Figure 3a). As SWAN is only a within-array probe type adjustment method it cannot remove the large technical variation present between the samples in the ageing+ dataset; therefore, we also applied SQN (43), which incorporates between-array normalisation. On the SQN pre-processed data, RUVm once again performed the best. Surprisingly, the performance of ISVA decreased substantially when the method was applied to data pre-processed with SQN (Figure 3b, Supplementary Figure 6e). The *limma* analysis including a factor for study performed better than ComBat on the SQN pre-processed data whilst the relative performance of the other methods remained the same (Figure 3b). Compared to SWAN, the use of SQN was of greatest benefit to the *limma* analyses (with and without adjustment for study) (Supplementary Figure 6a, b). The choice of pre-processing method had virtually no effect on RUVm (Supplementary Figure 6f) whilst the use of SQN was detrimental to ISVA (Supplementary Figure 6e).

**Figure 6.**
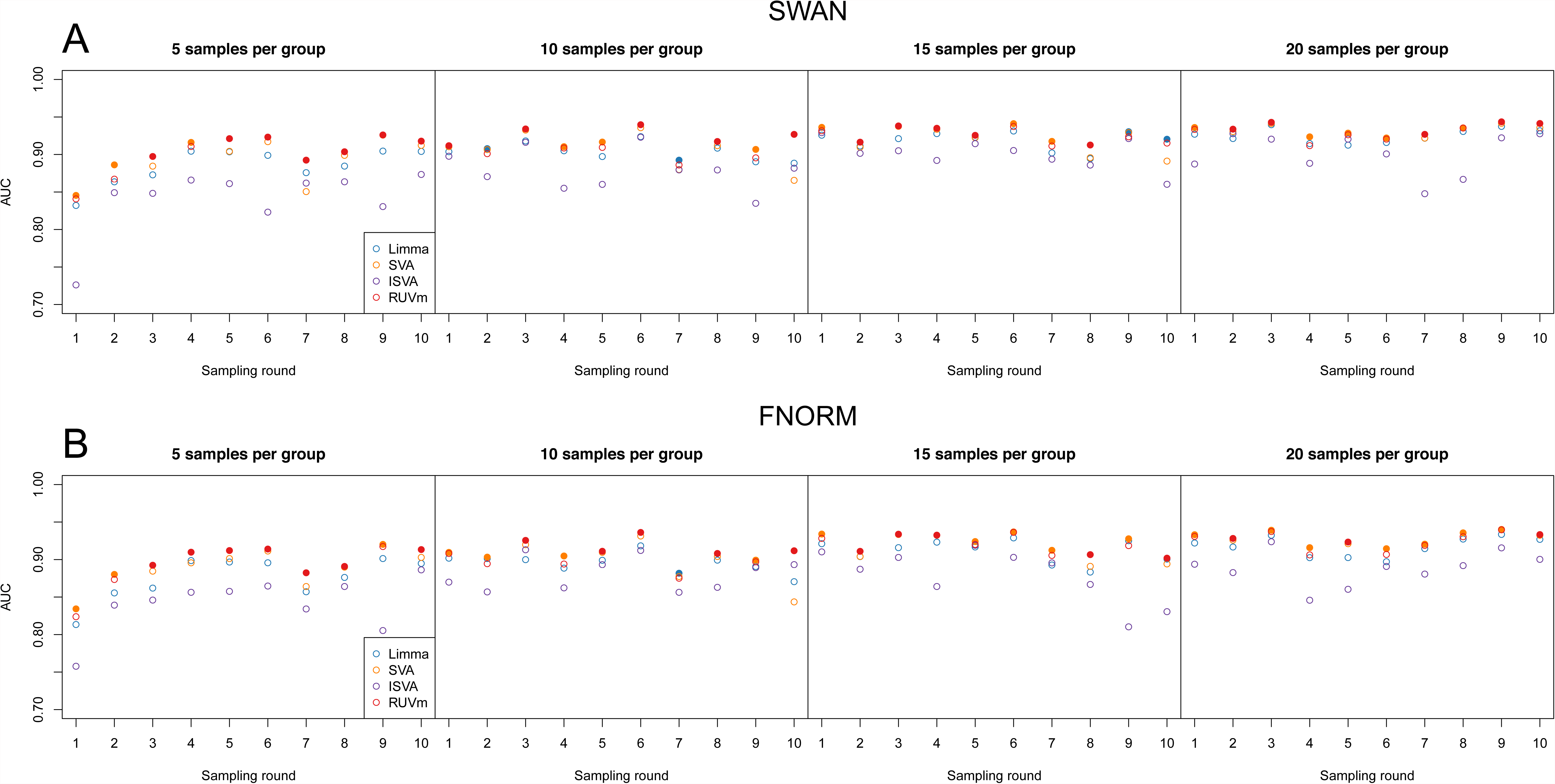
Performance of the various adjustment methods in DM analyses of smaller datasets subsampled from the TCGA LUAD data. The performance is expressed in terms of AUC. Each row represents the results for the same datasets pre-processed using either (A) SWAN or (B) FNORM. The individual panels in each row show the results at different levels of subsampling: 5, 10, 15 and 20 samples per group. The different colours correspond to the analysis methods used. A solid circle indicates that a particular method has the highest AUC for a single dataset.

### Smoking methylation data

Although the ageing+ data contains very large batch effects and other unwanted variation, the mean absolute difference in methylation between birth and later time points is in the order of ∼25%, which can be considered a relatively large effect size. Many EWAS aim to identify associations between methylation and diseases or environmental factors that have relatively small effects on the methylome (<10%). Unwanted variation can be a significant problem for such studies, particularly if the number of samples is not large. Consistent, but small, DNA methylation changes associated with smoking have been identified in multiple studies (46, 59–61). The differences in methylation between smokers and non-smokers are small (∼5%) and affect relatively few loci, genome-wide. Thus, the effect of smoking on methylation is a good example of a typical EWAS where unwanted variation in the data could be very problematic.

As such, we sought to evaluate the performance of RUVm and other adjustment methods in a DM analysis of 450k data of 200 current and 193 never smokers. The data was originally used to identify associations between methylation and rheumatoid arthritis by Liu et al. (6) and thus the samples came from both rheumatoid arthritis patients and controls, of both sexes and across a range of ages. The data was pre-processed with SWAN and filtered as described in the methods, leaving 398,313 probes. To assess performance we used a list of 187 CpGs identified by Zeilinger et al. (46) and subsequently replicated in a second cohort, as “true” positives (Supplementary Table 6). ECPs for use with RUVm were selected as the bottom 90% of probes from Stage 1.

Although the samples came from both rheumatoid arthritis patients and controls with mixed sex and age, the data did not appear to contain any large batch effects or other systematic technical variation (Supplementary Figure 7a). As there were no “known” batch effects we could only apply methods that did not require a batch to be specified. All of the analysis methods that incorporated an adjustment for unwanted variation (SVA, ISVA, RUVm) performed similarly, however, the unadjusted *limma* analysis was significantly less sensitive than the other approaches (Supplementary Figure 7b).

We then constructed a dataset with a large batch effect using a combination of samples from the Liu smoking data and samples from a second dataset, published by Hannum et al. (45). The Hannum dataset examined the association between methylation and ageing and did not provide information about smoking habits. However, as it consists of 656 samples, it is likely that they represent both smokers and non-smokers. To create the batch effect we randomly sampled 80 current smokers and 50 never smokers from the Liu data and then combined these with 70 random samples from the Hannum data, 20 of which were assigned as “smokers” and 50 as “never smokers” (Supplementary Table 3). This resulted in a dataset of 100 “smokers” and 100 “never smokers”, which will be referred to as the “smoking+” data (Figure 4a). This was intended to simulate two types of severe unwanted variation that may occur in an EWAS: a batch effect due to differences in sample collection and processing and unwanted variation due to misreported phenotypes and/or sample mix-ups.

We performed a DM analysis on the smoking+ data using the same methods applied to the ageing+ data and assessed performance using the 187 “true” positive CpGs previously described (46). ECPs for use with RUVm were once again selected as the bottom 90% of probes from Stage 1. Figure 5b shows that all the methods are significantly less sensitive on the smoking+ data than the Liu smoking data alone, as the unwanted variation is much more severe. However, RUVm is significantly more sensitive than the other adjustment methods on the smoking+ data (Figure 4b); RUVm manages to rank almost 100 of the 187 “true” smoking CpGs in the top 1000, whilst the other adjustment methods achieve ∼60 and the unadjusted *limma* analysis only manages ∼30.

We generated another two datasets using the same approach but with different randomly selected samples and different numbers of samples from the Liu and Hannum datasets (Supplementary Table 4, 5). RUVm, once again, performed better than the other methods in terms of sensitivity (Supplementary Figure 8), with a particularly marked improvement in the case with more extreme unwanted variation (Supplementary Figure 8a).

### Cancer methylation data

It has been observed that considerable heterogeneity exists in methylation within and between tumours (62). When performing a DM analysis of cancer versus normal samples the methylation data not only contains the expected methylation heterogeneity between tumours, it also contains unwanted biological variation arising from factors such as variable tumour purity due to presence of normal tissue, immune cells, etc. However, as cancer has a profound effect on the methylome the differences between cancer and normal can be very large (>50% of CpGs) and such comparisons also result in many more DM loci compared with other types of EWAS. Thus, unless a study has significant artefacts, cancer status is often the largest source of variation. Regardless of this, the presence of unwanted variation can affect how probes are ranked and whether they reach statistical significance, particularly if the sample size is not large. To investigate how the various adjustment methods perform in this scenario, we used the TCGA Lung Adenocarcinoma (LUAD) dataset. The “truth” was defined from an independent LUAD Agilent methyl-seq capture bisulfite sequencing dataset (see Methods).

We used all 31 of the TCGA LUAD normal samples, however, given that the complete LUAD dataset contains 427 tumour samples, we elected to use only the tumour samples that were assayed on the same chips as the normal samples, resulting in a more manageable number of 75 tumour samples. The 450k LUAD data was pre-processed using both SWAN (42) and *minfi* functional normalisation (FNORM), which is a between-array normalisation method for 450k data that is suitable for cancer (47). After separately filtering the TCGA LUAD 450k and methyl-seq datasets (see Methods), we intersected the remaining CpGs to identify the loci covered by both platforms, leaving 221,694 CpGs. RLE plots of the 450k data show that there is between-array variation that appears to be related to both cancer/normal status and chip (Figure 5). MDS plots of the 450k data also show evidence of clustering by chip and/or plate, particularly for the normal samples (Supplementary Figure 9a-d). Chip and plate effects can also be seen in the MDS plots of the 500 most variable INCs (Supplementary Figure 9e, f). An MDS plot of both the 450k and methyl-seq data using the 1000 most variable CpGs shows the separation between the cancer and normal samples and the relationship between the samples from different platforms (Figure 5). Using the methyl-seq samples, we defined 2,051 “true” positives (adj. p-value < 0.05) and 170,629 “true” negatives (adj. p-value > 0.9) by performing a DM analysis of the 5 tumour versus 5 normal methyl-seq samples using the Bayesian hierarchical model and Wald test approach implemented in the Bioconductor *DSS* package.

We then performed Stage 1 of the RUVm analysis. Examining the potential list of ECPs for use with RUVm shows that although the relationship between FDR cut-off and proportion of probes selected is still linear between 0.1 and 0.9, a high proportion (∼40%) of probes is significantly associated with cancer status at FDR adjusted p-value < 0.1 (Supplementary Figure 10). Consequently, the performance of RUVm is notably reduced when ≥80% of CpGs from the bottom of the list are designated as ECPs (Supplementary Figure 11c, d) but is consistent when they are selected based on FDR cut-off (0.1-0.9) as significantly cancer-associated probes are avoided (Supplementary Figure 11a, b).

We next compared the performance of four different methods in a DM analysis of the 450k LUAD data (31 normal versus 75 tumour samples); *limma*, SVA, ISVA and RUVm. RUVm was applied as previously described with the bottom 50% of probes from the Stage 1 ranked list designated as ECPs for Stage 2. An examination of the p-value histograms did not reveal a significant departure from the ideal for any of the methods (Supplementary Figure 12), although ISVA had visibly lower peaks near zero than any of the other methods (Supplementary Figure 12e, f). In an ROC analysis of the SWAN pre-processed data, *limma*, RUVm and SVA perform similarly. *Limma* performed only slightly better than RUVm, which in turn was marginally better than SVA. ISVA showed significantly reduced performance relative to the other methods (Supplementary Figure 13a). Using FNORM instead of SWAN pre-processing did not change the relative performance of the methods (Supplementary Figure 13b). Unexpectedly, apart from ISVA, all of the methods showed a slight decrease in performance with FNORM relative to SWAN (Supplementary Figure 14).

Although we did not utilise the entire TCGA LUAD 450k dataset, the 106 samples included in our analysis still provide a lot of power for detecting DM, as is evidenced by the competitive performance of *limma*, despite the significant amount of variability in the data. To investigate how the various analysis methods perform on a smaller number of samples we applied a subsampling strategy to the 106 samples used in the initial analysis. To retain approximately realistic chip effects, we first randomly selected chips from the 14 available chips (without replacement). The random selection of chips continued until the minimum number of samples required from each group was represented on the selected chips; for example, for an analysis of 5 cancer versus 5 normal samples, chips would be randomly sampled until there were at least 5 cancer and 5 normal samples present across the randomly selected chips. As this approach often resulted in more than the required number of samples, we then randomly sampled the exact number samples required from each group (e.g. 5) from only the samples present on the initially selected chips. We performed the subsampling 10 times for 5, 10, 15 and 20 samples per group, which produced 40 different datasets. This was done for both the SWAN and FNORM pre-processed data, resulting in 80 distinct datasets. A DM analysis was performed on each of the 80 datasets using the 4 methods previously mentioned.

Comparing the area under the curve (AUC) of the DM analyses of the subsampled data demonstrates that RUVm performed consistently well regardless of the number of samples and/or structure of the data (Figure 6). RUVm performed particularly well with only 5 samples per group, irrespective of the type of pre-processing used. SVA was also reasonably competitive, particularly with larger sample sizes and FNORM pre-processing; however, RUVm was computationally *much* faster. The AUC results are reflected in the ROC curves produced using the same data (Supplementary Figure 15).

## DISCUSSION

Due to their relatively low-cost per sample and single-nucleotide resolution coverage of >450,000 CpG sites across the human genome, Illumina’s 450k arrays are widely used in both small and large scale DM studies. However, despite the abundance of literature describing and evaluating methods for mitigating unwanted variation in the context of microarray gene expression studies there has been much less research focused on methylation data.

Here we present the novel approach, RUVm, which proposes the application of RUV-inverse in 2 stages (23) to 450k DM studies and show that it outperforms existing methods. In the first stage, we use the INCs for an initial DM analysis with RUV-inverse. In the second stage, we use the resulting rankings of the CpG probes to select ECPs for a second DM analysis with RUV-inverse. As the INCs capture various types of technical variation such as plate or chip (Supplementary Figure 10, 16), their use in Stage 1 is often an improvement over an unadjusted analysis. However, caution is advised if the INCs are affected by unwanted variation that is strongly correlated with the factor of interest as this is likely to be detrimental to the performance of RUVm. Cases displaying strong correlation between technical variation and the factor of interest are often reflective of poor experimental design and the results of any method must be treated with scepticism.

An important consideration for the application of RUVm is selection of ECPs. ECPs need to be selected on a case by case basis and the results should be carefully evaluated. We investigated the effect of ECP selection on the overall performance of RUVm for both the ageing+ and LUAD cancer data. We considered selection of ECPs based on a series of FDR cut-offs as well as selecting a fixed percentage of probes from the bottom of the ranked list. Our investigation showed that examining the results of the DM analysis performed in Stage 1 provides a reasonable indication of the final amount of DM and is thus a valuable guide to selecting ECPs. Supplementary Figures 3 and 10 highlight the difference between the ageing+ and cancer studies, respectively, in terms of DM and the acceptable number of probes that ECPs should be drawn from. Fewer DM CpGs are expected to be found in the ageing+ data compared with the cancer data and, as expected, more probes can be designated as ECPs in the ageing+ study compared to the cancer study, without detriment to performance. Based on our analyses, we suggest that selecting ECPs based on an FDR cut-off around 0.5 or approximately the bottom 50% of the ranked list should work well in the absence of other information.

Although the use of ECPs is likely to be a good option for many methylation studies, they are not the only option for using RUVm to analyse methylation data. Recalling that the criteria for selecting a good set of negative controls are (a) that they should not be truly associated with the factor of interest and (b) that they should be affected by unwanted variation, one can take a number of approaches to choosing negative controls. For example, a researcher may know *a priori* that, for their particular experiment, there is a set of CpGs that should be invariant with respect to the effect they are investigating and as such could be useful negative controls. Alternatively, a set of negative controls could be defined using data from previous experiments, as in the study by Jaffe and Irizarry (8). Whatever strategy is employed, it is always prudent to evaluate how an adjustment is performing; examples of some useful approaches can be found in this study and are also discussed in detail in Gagnon-Bartsch and Speed (22) and Gagnon-Bartsch et al. (23).

We have demonstrated that the RUVm can correct for various types of unwanted variation in a DM analysis of 450k data. In particular, we have found that RUVm consistently outperforms other methods when the effect of unwanted variation in the data is larger than the factor of interest. Specifically, using the ageing+ dataset, which was dominated by a very large cell type batch effect and cell-type heterogeneity issues, we showed that RUVm performed better than other methods irrespective of the type of pre-processing applied to data. With a more subtle phenotype, such as smoking, all methods that adjusted for unwanted variation performed similarly on data with no discernible batch effects (Figure 5). However, on the smoking+ data, which incorporated a large a batch effect and several misallocated samples, RUVm ranked almost twice as many true positives in its top 1000 CpGs than any of the other adjustment methods. Consequently, we expect that RUVm should be particularly beneficial for the analysis of very “messy” datasets such as those that seek to combine samples from multiple labs/studies.

Cell-type composition has also been demonstrated to be an issue in methylation studies (11–15). For the analysis of the ageing+ data, to ensure that the methods we were evaluating were identifying CpGs with real age-associations, we carefully assessed the ageing “truth” data to ensure that cell type composition changes were not significantly confounded with age. We also adjusted for cell type proportion estimates when defining “true” positives and negatives. We thus expected that, in addition to accounting for the batch effect, methods that were able to adjust for cell type composition should rank “truly” age-associated CpGs more highly than cell type associations. RUVm was able to recover almost 80% of “true” positives irrespective of the pre-processing method with a relatively low false positive rate. RUVm also performed more consistently than other methods when comparing normal tissue and solid tumours, which are often very heterogeneous, particularly when the sample size was low. We propose that this demonstrates that RUVm is able to adjust for fluctuations in cell type composition, enabling it to prioritise true associations over spurious effects. Jaffe and Irizarry (8) made a similar observation in their study when they applied RUV-2 (22). Nevertheless, we would still advise that any methylation study of a mixed-cell population incorporate rigorous checking for serious confounding between cell type composition and the factor of interest.

The RUVm approach proposed herein is a versatile method that effectively accounts for unwanted variation in DM analyses of 450k data across a wide variety of studies. RUVm performs particularly well in situations where the sources of unwanted variation are large relative to the factor of interest and therefore we believe this method will be of great utility to the EWAS community. All of the core RUV methods, including RUV-inverse, have been implemented in R and are available in the CRAN package *ruv* (http://cran.r-project.org/web/packages/ruv/index.html). Functions extending the *ruv* package to align it with the *limma* framework and for its application to 450k data have been implemented in the *missMethyl* Bioconductor package (http://www.bioconductor.org/packages/release/bioc/html/missMethyl.html).

## ACKNOWLEDGEMENT FUNDING

This work was supported by the National Health and Medical Research Council, Project Grant APP1051402 to AO, Australia Fellowship to TPS, and Career Development Fellowship to AO. Funding for open access charge: National Health and Medical Research Council.

---

Supplementary Table 1. Ageing samples used in this study. Supplementary Table 2. Liu et al. (6) smoking samples used in this study.

Supplementary Table 3. Hannum et al. (45) and Liu et al. (6) samples used in this study to create the smoking+ dataset (Combination 1).

Supplementary Table 4. Hannum et al. (45) and Liu et al. (6) samples used in this study (Combination 2).

Supplementary Table 5. Hannum et al. (45) and Liu et al. (6) samples used in this study (Combination 3).

Supplementary Table 6. 187 smoking CpGs identified and replicated by Zeilinger et al. (46) used as true positives in this study.

Supplementary Table 7. TCGA LUAD samples used in this study. Supplementary Table 7. TCGA LUAD samples used in this study.

